# Antifungal resistance mechanisms and nosocomial transmission of *Nakaseomyces glabratus*: genomic investigation and observational study in Melbourne, Australia

**DOI:** 10.64898/2026.06.22.733717

**Authors:** Andrew Gador-Whyte, Torsten Seemann, Louise Judd, Kristy Horan, Jake Lacey, Ana Traven, Diane Daniel, Romain Guérillot, Stefano Giulieri, Sara Vogrin, Maria Duarte Aguilera, Marcel Leroi, Gemma Reynolds, Benjamin P Howden, Norelle Sherry, Jason Kwong

## Abstract

*Nakaseomyces glabratus* (*Candida glabrata*) is a WHO high-priority fungal pathogen associated with fungal antimicrobial resistance (fAMR). Given nosocomial transmission occurs sporadically, resistant strains could be transmitted, a concern for critically ill patients. We conducted a genomic investigation and retrospective observational study of *N. glabratus* to identify any nosocomial transmission of fAMR and understand resistance mechanisms and clinical and demiological factors among patients at a quaternary hospital in Melbourne, Australia. We selected stored *N. glabratus* with and without fAMR associated with similar patient clinical characteristics and performed whole genome sequencing. Clinical and epidemiological data were extracted from medical records. Phylogenetic, mutational, copy-number variation (CNV) and mitochondrial genomic analyses were performed, with a focus on the fAMR gene *PDR1*. Of 54 isolates collected over seven years, 20 (37%) were fluconazole-resistant and four (7%) had elevated flucytosine minimum inhibitory concentrations (MICs) (range 2-32 μg/ml). There were no significant clinical differences between patients with and without fluconazole resistance. Most (55%) fluconazole-resistant isolates carried *PDR1* mutations. Resistance was distributed throughout the phylogeny suggesting predominantly independent acquisition. However, a cluster of four resistant isolates with the same *PDR1* mutation suggested nosocomial transmission. One probable *ERG11* gene duplication, and two petite variants with apparent mitochondrial genomic deletions, were seen in association with fluconazole resistance. In this study, we identified a small probable nosocomial fAMR transmission cluster, and novel variants in *PDR1*, *ERG11* and *FCY2* associated with fAMR phenotypes. Future study should confirm functional impacts and systematically investigate for nosocomial transmission of resistance, including colonisation states.

## INTRODUCTION

*Nakaseomyces glabratus* (formerly *Candida glabrata)* is a human-adapted pathogenic yeast which is second only to *Candida albicans* as a cause of fungal bloodstream infections and other invasive infections (1). Invasive infections carry a high mortality rate, especially in critically ill and immunocompromised patients (2). *N. glabratus* is recognised as a High Priority organism in the 2022 WHO Fungal Priority Pathogen list (3). Previous studies have demonstrated sporadic nosocomial transmission (4, 5). *N. glabratus* possesses a haploid and plastic genome that facilitates the emergence of fungal antimicrobial resistance (fAMR) (6).

*N. glabratus* is intrinsically less susceptible to the first-line triazole antifungal, fluconazole. The triazoles, which inhibit synthesis of the membrane lipid ergosterol, are widely used for treating *N. glabratus* infections (2). The echinocandins (such as caspofungin) inhibit synthesis of the cell wall component 1,3-b-D-glucan, and are the first-line treatment for *N. glabratus* bloodstream infections (7). Flucytosine, an inhibitor of nucleic acid synthesis, has applications in urinary tract infections (7). The polyene drug amphotericin B, which binds to ergosterol causing membrane instability, is sometimes used in complex infections(8).

Fluconazole resistance occurs in up to 48% of isolates (2). Acquired resistance is usually due to enhanced efflux, usually from gain-of-function mutations in the transcription factor gene *PDR1* (9–11). Other mechanisms include target over-expression due to *ERG11* copy-number variation (CNV) (12) and mitochondrial genome deletions leading to a resistant phenotype mediated by *PDR1* overexpression and manifesting as petite variants (11, 13, 14) Echinocandin resistance is uncommon (generally <5%) (2) and usually due to mutations in ‘hot spot’ regions of the glucan synthase gene *FKS2* (1, 8). Flucytosine reduced susceptibility appears rare (reported range 0-4.7%), and is associated with mutations in *FCY1, FCY2* or *FUR1* (2, 8). Polyene non-susceptibility, although generally <2% (2), may arise due to loss-of-function mutations in ergosterol pathway genes including *ERG11, ERG6, ERG2,* or *ERG3* (8).

Understanding how frequently nosocomial transmission of fAMR occurs, and the epidemiological and pathogen factors underpinning it, is critical to prevention in a vulnerable hospitalised patient cohort. Genomic sequencing has the potential to enable timely detection of fAMR transmission and elucidation of resistance mechanisms. Interrogating potential emergence and transmission of fAMR within healthcare requires granular epidemiological data, including relating to antifungal prescribing and bed locations. Here, we report results of a retrospective observational study and genomic investigation of *N. glabratus* infections among hospitalised patients in Melbourne, Australia. We sought to identify and understand the mechanisms of, and clinical and epidemiological factors associated with, fAMR and identify any nosocomial clonal transmission of fAMR.

## METHODS

### Setting and antifungal susceptibility testing

This study was conducted at a quaternary hospital in Melbourne, Australia (approximately 1000 beds) which includes specialty referral services including liver transplant and a 30-bed Intensive Care Unit (ICU).

### Isolate selection and clinical data collection

We included clinical isolates from 2018 to 2024 exhibiting phenotypic non-susceptibility to commonly used antifungal medications (echinocandins, triazoles, flucytosine and amphotericin B). For comparison, we selected susceptible isolates derived from patients with a similar range of clinical infections (Figure 1). For simplicity, we used the following definition for “Non-susceptible” in this study: (i) resistance to fluconazole or echinocandins by the Clinical and Laboratory Standards Institute (CLSI) breakpoints; (ii) susceptibility to fluconazole but with minimum inhibitory concentrations (MICs) at least 4-fold higher than the CLSI epidemiological cutoff value (ECV) for two other triazoles; or (iii) flucytosine MIC ≥ 0.5 µg/ml. “Susceptible” was defined as susceptible, dose-dependent (SDD) to fluconazole, susceptible to echinocandins and not meeting criteria for Non-susceptibility. Further details on isolate selection and these definitions (including selection of a cutoff for flucytosine) are provided in the Supplemental Material. Demographics, clinical and treatment characteristics and bed location data were obtained from the electronic medical record.

**FIGURE 1.**
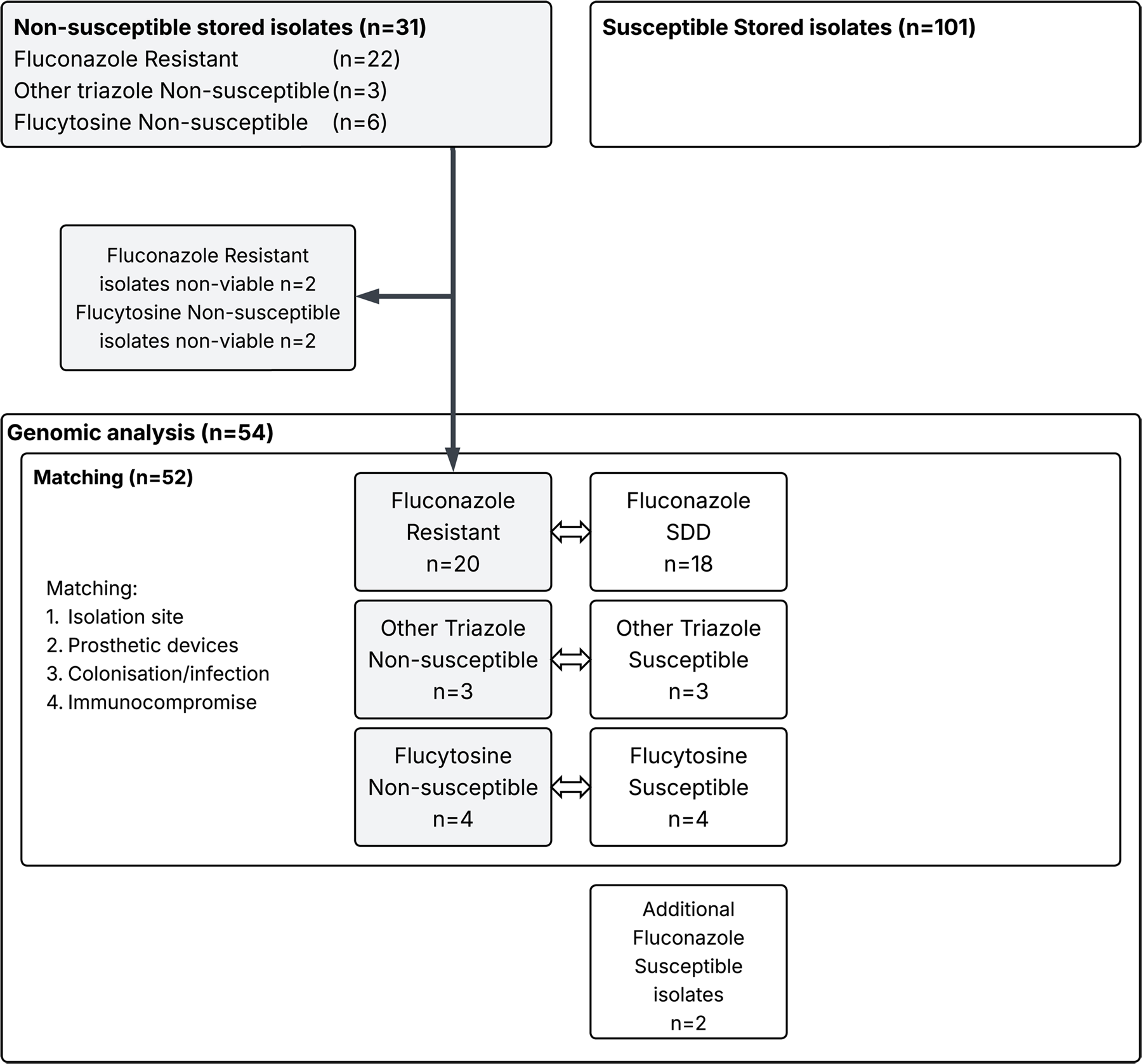
Isolate selection schema including patient matching to achieve a balanced population for epidemiological and genomic comparison. Abbreviations: SDD: Susceptible, Dose-Dependent

### Culture, quality control and confirmation of susceptibility profiles

*N. glabratus* isolates stored at -80°C were cultured on Sabouraud dextrose agar with gentamicin (ThermoFisher, Australia) in air at 35-37°C for 24-48h. Species identification was performed using MALDI-TOF MS (VITEK® MS PRIME, Biomérieux, France) and broth microdilution performed using Sensititre™ YeastOne™.

MICs were interpreted according to CLSI breakpoints (M27M44S-Ed3 (2022)) (15) or epidemiological cutoff values (ECVs) (M57SE-Ed4 (2022)) (16). Essential agreement (EA) was defined as MICs from original and repeated testing falling within 2 doubling dilutions, as per CLSI. Categorical agreement (CA) was defined as concordance in susceptibility category between original and repeated testing. Isolated caspofungin discrepancies were not reported because of known major errors with caspofungin testing (15).

### Whole genome sequencing

Isolates underwent Illumina whole genome sequencing as per local research laboratory protocols (see Supplemental Material for details).

### Statistical analysis

Statistical analysis was performed in RStudio (17). Associations between continuous nonparametric variables and fluconazole susceptibility were assessed with the Mann-Whitney U test, and between categorical variables using the Chi-squared or Fisher’s exact test, as appropriate. MIC distribution bar plots were generated using ggplot2 v3.5.1 (18). Table S10 summarises the R software packages used for statistical analysis and figures.

### Bioinformatic analysis

Sequence read quality control was performed using seqkit v2.1.0 and kraken2 v2.1.2 to ensure a minimum average read depth (40x), GC content within 2% of expected and correct species identification and minimal (<1%) contamination by *k*-mer classification. Two variant calling pipelines were used : (i) Bohra v2.3.7 (19); and (ii) the Genome Analysis Toolkit (GATK) v4.6.2.0 (20). Mutations in genes with possible fAMR associations (see Table S1) were annotated using bcftools csq (21) and assessed against the FungAMR and AFRbase databases, and previous studies (5, 22–24). Contigs were also analysed for FungAMR mutations using ChroQueTaS v0.6.0 (23). For MLST, single-locus variants (SLV) were represented provisionally as SLVs of the closest MLST. Phylogenetic tree visualisation and annotation was performed in RStudio using ggtree v3.14.0 and other packages (see Supplemental Material) (25).

Phylogenetically related isolate pairs with differing phenotypic antifungal susceptibility were assessed for novel mechanisms. Copy-number variation (CNV) was explored by assessing the average read depth of individual genes and chromosomes against the median depth with samtools coverage (26). Putative mitochondrial genome deletions were ascertained by (i) visual inspection of read alignment to the mitochondrial genome with Tablet (27), (ii) estimating coverage across seven mitochondrial genes with samtools coverage, and (iii) *de novo* assembly of reads mapping to a mitochondrial genome reference, assessing assembly size and completeness including with Bandage (28). Further methodological details are included in the Supplemental Material.

To place our sequences in a global context, paired Illumina sequence reads were downloaded from the NCBI Sequence Read Archive (SRA) (29) and visualised using mashtree (30) with *PDR1* mutations detected using Ariba (31).

Full bioinformatic methods are provided in the Supplemental Material.

## RESULTS

### Isolate selection

Isolate selection is outlined in Figure 1. Suitable matched Susceptible isolates were available for 25 Non-susceptible isolates. Two additional Susceptible isolates were included to provide a balanced population for phylogenomic and mutational analysis (27 Non-susceptible, 27 Susceptible). Sample collection site and year are provided in Table S3.

### Antifungal susceptibility testing

Antifungal susceptibility data is provided in Table S4. One fluconazole-resistant isolate demonstrated low-level anidulafungin resistance (MIC 0.5 μg/ml) but susceptibility to micafungin (0.03 μg/ml), and Non-susceptibility to amphotericin B (MIC 4 μg/ml). Among Susceptible isolates, most MICs were above the ECV for voriconazole (50/54, 92.6%). All but four had low (<=0.06 μg/ml) flucytosine MICs.

Essential non-agreement between original and repeated testing was seen in four isolates (lower MIC on repeat testing in all cases). Three fluconazole-resistant isolates fell into the SDD (<=32 μg/ml) range on repeat testing. For other triazoles, as most isolates had had MICs close to the ECV, categorical non-agreement was frequent (25/54, 46%) despite high EA (50/54, 93%). The anidulafungin-resistant and amphotericin B Non-susceptible isolate tested intermediate to anidulafungin (0.25 μg/ml) and wild-type to amphotericin B (1 μg/ml) on repeat testing.

### Clinical characteristics

Isolates were predominantly from urine (19/53, 35%) and blood culture (18/53, 35%) samples, followed by intraabdominal (12/53, 22%), thoracic (2/53, 4%), and other sites (2/53, 4%) (See Table S3). Clinical characteristics were similar between the groups with and without fluconazole resistance (see Table 1). In the 12 months preceding isolate collection, 15/54 (28%) of patients had exposure to systemic antifungals (triazoles 11/54, 20%; echinocandins 4/54, 7%). Exposure to any triazole was non-significantly more frequent in the fluconazole-resistant group (5/20, 25%; vs 5/34, 15%). Triazole exposure ranged from 7 to 27 days. The patient with a low-level anidulafungin resistant isolate had 13 days of recent echinocandin exposure in the setting of liver transplantation.

**Table 1.** Comparison of clinical, treatment and epidemiological characteristics of patients with an included *N. glabratus* isolate.

| Variable | Frequency, fluconazole-resistant group<br>n (%) | Frequency, fluconazole-susceptible group<br>n (%) | p-value |
| --- | --- | --- | --- |
| Diabetes | 7 (35%) | 16 (47%) | 0.56 |
| Any triazole exposure (12 months) | 5 (25%) | 5 (15%) | 0.56 |
| Urinary tract foreign body | 5 (25%) | 6 (18%) | 0.77 |
| Any immunocompromising condition | 7 (35%) | 17 (50%) | 0.43 |
| - Solid organ transplant | 1 (5%) | 6 (18%) | 0.24 |
| - Haematological malignancy | 1 (5%) | 4 (12%) | 0.64 |
| - Solid organ malignancy | 3 (15%) | 5 (15%) | 1.0 |
| - Inflammatory bowel disease | 1 (5%) | 1 (2.9%) | 1.0 |
| - Rheumatological disorder | 2 (10%) | 2 (5.9%) | 0.62 |
| Spinal cord injury | 2 (10%) | 1 (2.9%) | 0.33 |
| Abdominal surgery (12 months) | 5 (25%) | 8 (24%) | 1.0 |
| Total parenteral nutrition (12 months) | 6 (30%) | 3 (8.8%) | 0.062 |
| Bacteraemia (12 months) | 4 (20%) | 6 (18%) | 1.0 |
| Dialysis (3 months) | 3 (15%) | 4 (12%) | 1.0 |
| Central venous catheter (3 months) | 11 (55%) | 15 (44%) | 0.62 |
| Indwelling urinary catheter (3 months) | 10 (50%) | 20 (59%) | 0.73 |
| Mechanical ventilation (48 hours) | 3 (15%) | 6 (18%) | 1.0 |
| ICU admission (current or past 48 hours) | 3 (15%) | 10 (29%) | 0.33 |
| *Candida* colonisation (2 weeks) | 3 (15%) | 7 (21%) | 0.73 |
| Variable | Duration (days), fluconazole-resistant group (median, IQR) | Duration (days), fluconazole-susceptible group (median, IQR) | p-value |
| Inpatient ward admission | 22 (9-42.25) | 12 (4-32.25) | 0.25 |
| ICU admission | 1 (0-4) | 0 (0-6.25) | 0.75 |

**Abbreviations:** ICU: Intensive care unit

### SNP analysis, phylogenetic analysis, sequence typing and genomic epidemiology

Sequence quality was adequate for all isolates. There was close concordance in fAMR mutations between Bohra and GATK using bcftools csq, and ChroQueTas using the custom database (See Supplemental Material). Pairwise SNP distances ranged between 202 and 63913 SNPs. The isolate collection formed a polyphyletic structure (see Figure 2). The most frequent MLSTs were ST26 (n=13), ST10 (n=6), ST7 (n=6) and ST3 (n=6). Within MLST groups, there was no progressive increase in Non-susceptibility over time, suggesting fAMR was more likely to arise sporadically (see Figure 3).

**FIGURE 2.**
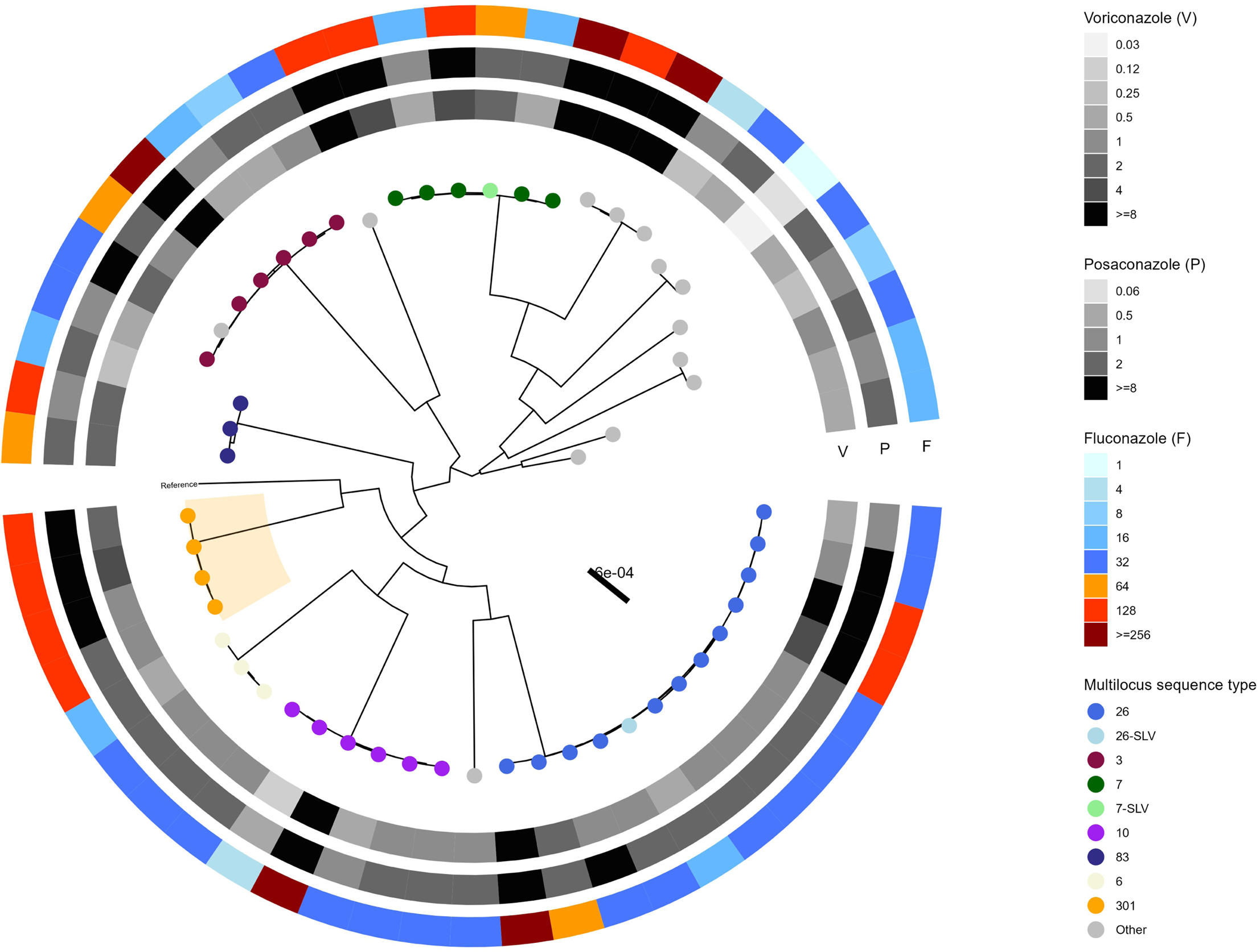
Maximum likelihood phylogenetic tree annotated by triazole susceptibility (heatmaps) and sequence types (tip points). Cluster of ST301 isolates highlighted in orange. **Abbreviations**: SLV: Single locus variant (single nucleotide difference in one allele separating from nearest MLST – provisional designation).

**FIGURE 3.**
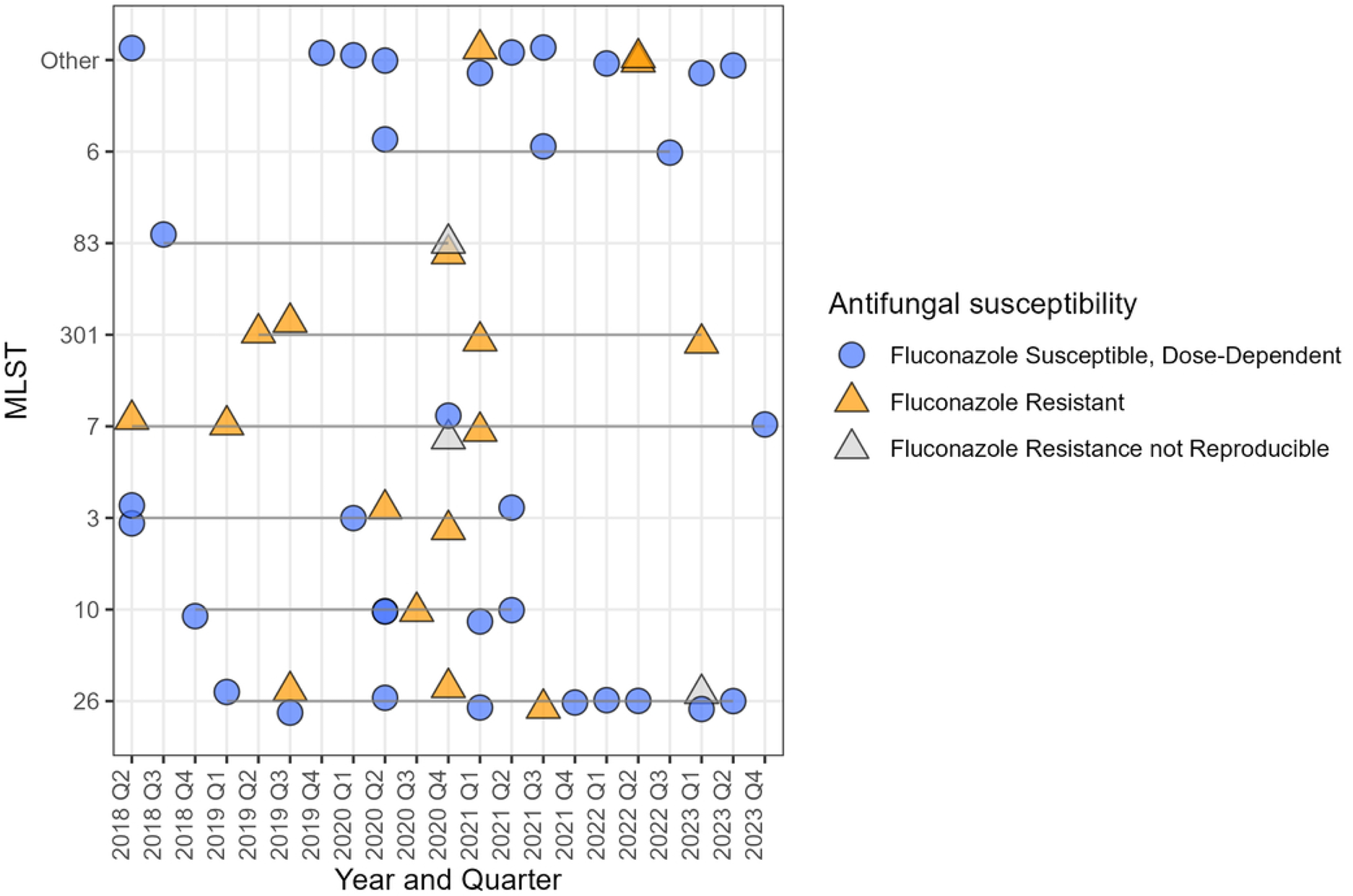
Timeline of isolates by multilocus sequence typing and antimicrobial resistance phenotype.

The fluconazole-resistant ST301 cluster (n=4) included four patients with prior ICU admission. Three had surgical conditions. All isolates harboured the same *PDR1* mutation, M957I, seen only in this cluster, but previously described in association with fluconazole resistance (9). Isolate pairs in this cluster were separated by 256 to 417 SNPs.

Study sequences were visualised in the global context with 1127 short-read whole genome sequences downloaded from NCBI SRA, confirming monophyly of the ST301 cluster (See Supplemental Material and Figure S1).

### Mutations in fAMR genes

Most (11/20, 55%) fluconazole-resistant isolates harboured mutations in *PDR1* that were restricted to resistant isolates. Efflux pump gene mutations were frequent (13/20, 65%) (Table 2). No resistance mechanism was identified in 10 Non-susceptible isolates, including all three with high MICs to two or more triazoles but susceptible to fluconazole (Table 2). Most *PDR1* mutations were previously described; novel *PDR1* mutations included L833H in a fluconazole-resistant isolate (same position as L833V listed in FungAMR) (8); and a frameshift at position 179 in one isolate, likely explaining its triazole-hypersusceptible phenotype (fluconazole and voriconazole MICs 1.0 and 0.03 μg/ml respectively) (Table 3) due to a presumed loss of *PDR1* function. Viewing the *PDR1* mutations occurring at each fluconazole MIC level identified (a) a number of mutations relative to the reference which are not fAMR-associated; (b) a higher burden of *PDR1* mutations among the fluconazole-resistant isolates; (c) a multiplicity of mutations associated with a common phenotype; and (d) a mutation associated with fluconazole resistance in clustered isolates, M957I (Figure 4) (9). Most (9/11, 82%) of the *PDR1* mutations occurring in resistant isolates only were in three previously described transcriptionally-active functional domains (9), including all such mutations with ≥90% mapped read support (see Figure 5).

**FIGURE 4.**
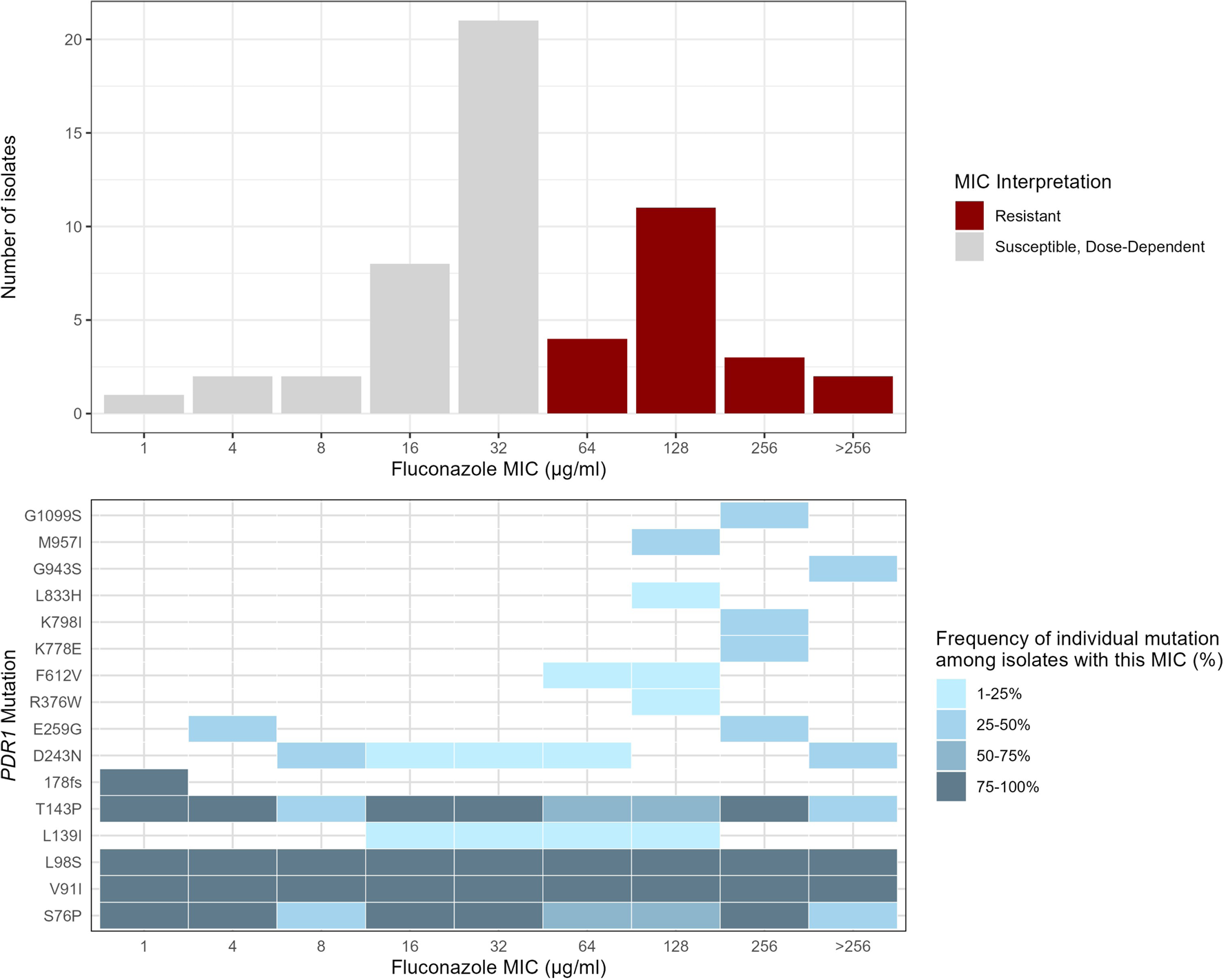
Fluconazole MICs with heatmap of frequency of PDR1 mutations by MIC level. <u>Upper chart</u>: number of isolates with each fluconazole MIC, coloured by categorical interpretation (Resistant or Susceptible, Dose-Dependent according to CLSI M59). <u>Lower chart</u>: frequency heatmap of PDR1 mutations occurring in isolates at each MIC level. Tiles shaded according to the proportion of isolates with a given MIC that demonstrated this mutation. MIC, minimum inhibitory concentration.

**FIGURE 5.**
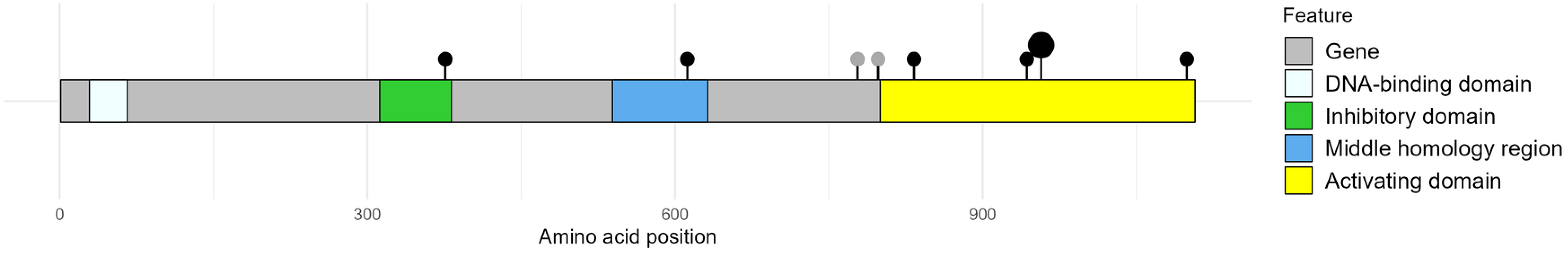
Plot of location of PDR1 amino acid mutations overlaid on the linear structure of gene product with four key functional domains highlighted. Functional domains represented by coloured bands. Amino acid position of mutations occurring only in fluconazole-resistant isolates represented by dots (black: ≥90% reads mapping to variant, grey: <90% reads mapping to variant). Amino acid coordinates of functional domains: DNA-binding domain, 20-66; Inhibitory domain, 312-382; Middle homology domain, 539-632; Activating domain, 800-1107.

**Table 2.**
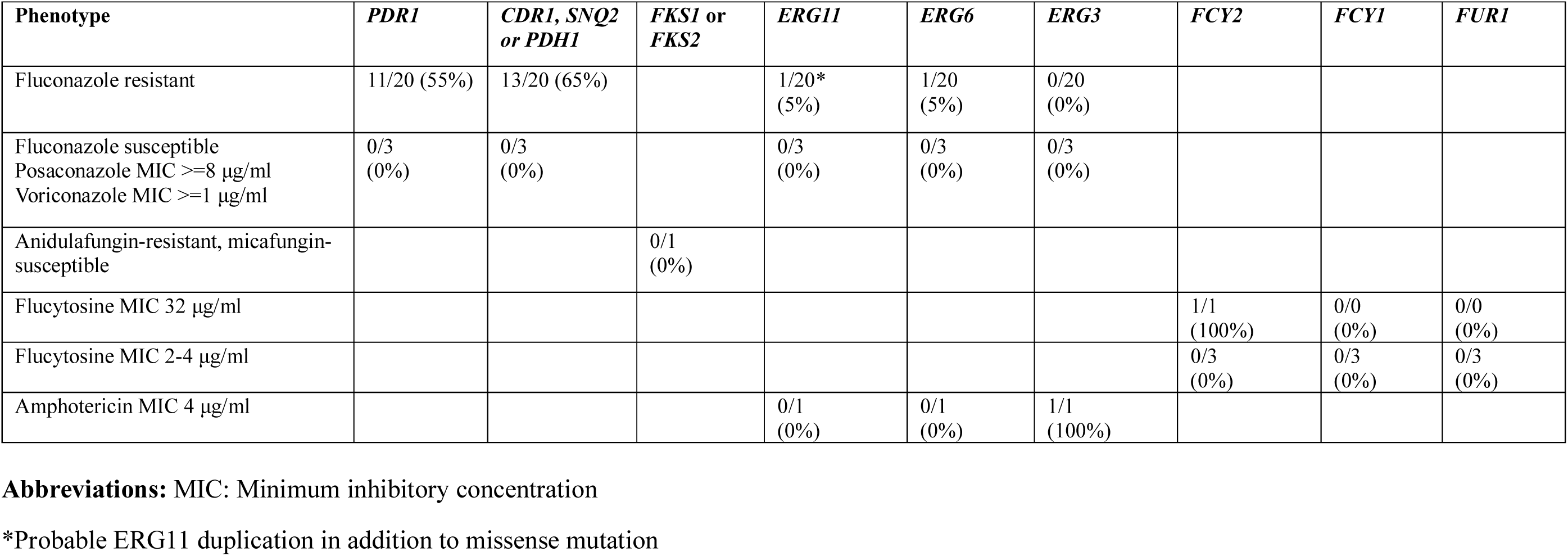
Frequency of mutations in genes associated with fAMR occurring only in Non-susceptible isolates, by fAMR phenotype.

**Table 3.** Mutations in fAMR -associated genes occurring only in Non-susceptible isolates, by fAMR phenotype: previously reported and novel mutations.

| Gene | Gene product | fAMR phenotype observed | Reported impacts | Mutation: previously reported | Novel mutation |
| --- | --- | --- | --- | --- | --- |
| ***PDR1*** | Pleiotropic drug response element (zinc finger transcription factor) | Fluconazole resistance | Triazole (gain-of-function) | R376W<sup>(23, 24)</sup><br>K778E*<sup>(23)</sup><br>G943D<sup>(22, 23)</sup><br>G943S<sup>(23, 24)</sup><br>G1099S<sup>(23, 43)</sup><br>F612V<sup>(23)</sup><br>M957I<sup>(23)</sup> | L833H†<br>K798I* |
| ***CDR1*** | ABC-type multidrug transporter | Fluconazole resistance | Triazole (uncertain) | F530S<sup>(22)</sup><br>I1478M<sup>(22)</sup> | V228L<br>F551S ‡<br>S841L<br>G881S |
| ***PDH1***<br>**syn.:**<br>***CDR2*,**<br>***PDR15*** | ABC-type multidrug transporter | Fluconazole resistance | Triazole (uncertain) | V734I<sup>(34)</sup> | L1451M |
| ***SNQ2*** | ABC-type multidrug transporter | Fluconazole resistance | Triazole (uncertain) | G1421S<sup>(34)</sup> | K351E<br>1421-fs |
| ***FCY2*** | Cytosine permease | High flucytosine MIC | Flucytosine (putative loss-of-function) |  | 341-fs (complex) |
| ***FCY21*** | Purine-cytosine transporter | High flucytosine MIC | Flucytosine (putative loss-of-function) |  | W291L<br>L323*<br>W502* |
| ***ERG11*** | Lanosterol 14α demethylase | Fluconazole resistance | Triazole (target binding site alteration)<br>Polyene (loss-of-function) | | I91V |
| ***ERG6*** | Sterol C-24 methyl-transferase | Fluconazole resistance | Polyene (loss-of-function) |  | V335I |
| ***ERG3*** | Sterol C-5 desaturase | Amphotericin B non-wildtype | Triazole (loss-of-function)<br>Polyene (loss-of-function) |  | H218R |

For the efflux pump genes *CDR1*, *PDH1* and *SNQ2*, the best-documented mechanism of resistance is overexpression; missense mutations in these genes might theoretically alter affinity of drug binding to efflux pumps but require functional confirmation and are therefore listed as of ‘uncertain impact (column 4) pending functional confirmation. fs= frameshift; syn=gene synonyms; \*supported by <90% of reads (total read depth at this site >90). fAMR= fungal antimicrobial resistance.
† Same position as L833V recorded in FungAMR
‡ Same position as F551L reported in De Luca et al, 2026 (34)

The isolate with flucytosine MIC 32 μg/ml had a predicted frameshift in *FCY2* due to a single-nucleotide insertion and 3 SNPs. This and another Non-susceptible isolate (MIC 4 μg/ml) carried stop mutations (W502* and L323*) in *FCY21*, a purine-cytosine permease gene with sequence similarity to *FCY2* (32). The isolate with discordant echinocandin susceptibility findings and Non-susceptibility to amphotericin B carried mutations in *FKS2* (T926P), *FKS1* (G14S) and *FKS3* (A42V and T1676S) and an *ERG3* mutation, H218R, that has not previously been identified in amphotericin B-non-susceptible isolates.

### Copy-number variation

Probable CNV was identified in isolate with maximal MICs to all triazoles in a patient without recent antifungal exposure (Figure 6). This was an apparent *ERG11* duplication due to a Chromosome E duplication. This occurred concurrently with an *ERG11* mutation, I91V, not previously linked to triazole susceptibility, and a probable resistance mutation in *PDR1*, G943S. Extending the exploratory CNV analysis to the global dataset, putative chromosome E duplications occurred in 13 isolates in an earlier Australian study (22) (fluconazole-susceptible, n=11; resistant, n=2) and eight international isolates (resistant, n=2, susceptible, n=2, susceptibility not provided, n=4).

**FIGURE 6.**
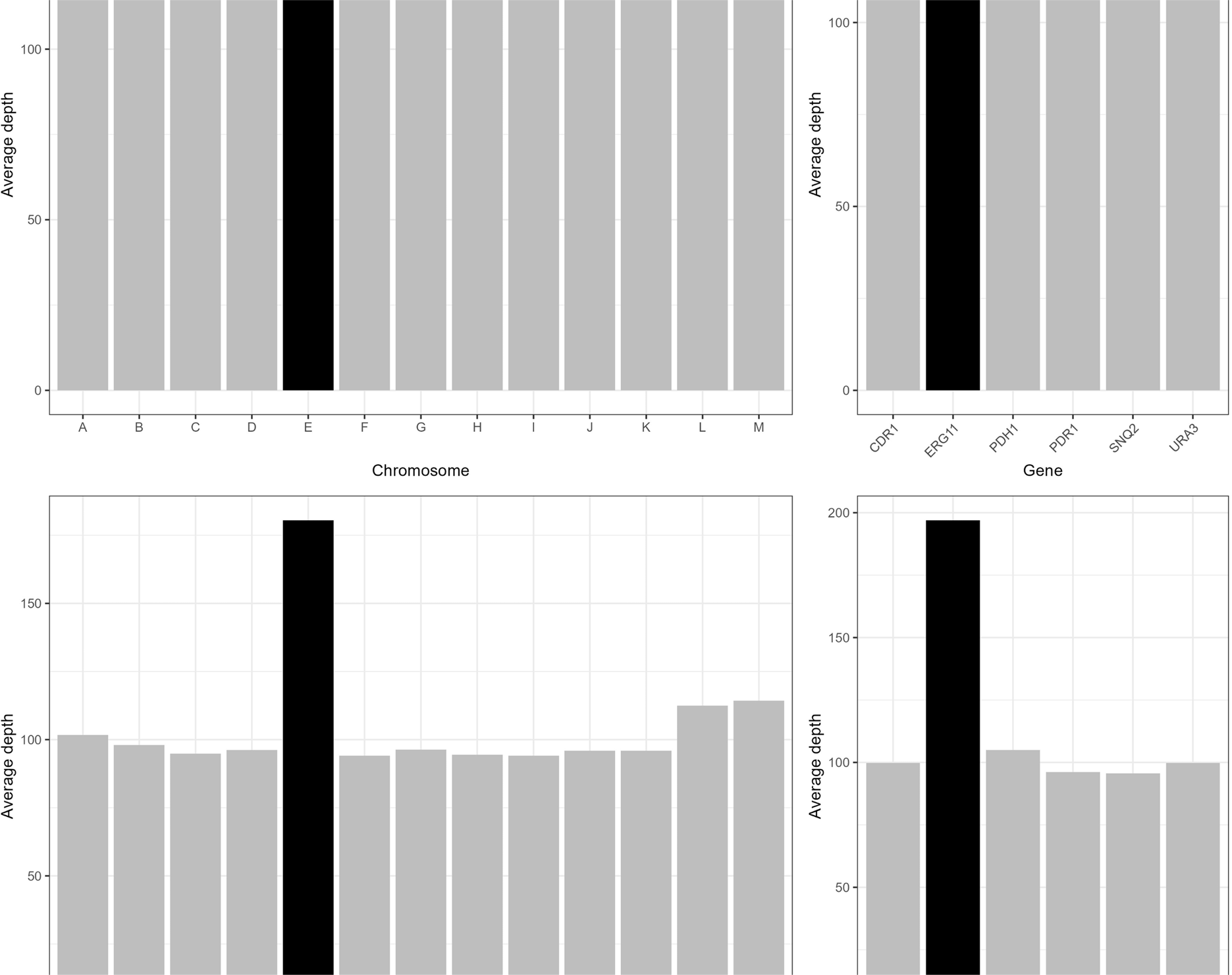
Copy-number variation by chromosome and gene illustrated using read depth data (number of times region covered by sequence reads) from samtools coverage. <u>Upper panels</u>: median average depth by chromosome (left) and gene of interest (right), all isolates. <u>Lower panels</u>: median average depth by chromosome (left) and gene of interest (right), isolate with apparent chromosome E duplication leading to ERG11 duplication. Chromosome E and ERG11 gene highlighted (black).

### Mitochondrial genome analysis

Most isolates had high coverage across seven mitochondrial genes, apart from two (AUH_NAGL1 and AUH_NAGL9) with no coverage across the majority of the genes (See Table S6), suggesting that they have lost the mitochondrial genome, with reads visually mapping only to a small section (approximately 700 and 1000 bp respectively) at the beginning of the mitochondrial genome. *De novo* assembly produced 1-3 contigs and a maximum contig length approximating 20 000 bp, the length of the mitochondrial genome (33), in all isolates except AUH_NAGL1 and AUH_NAGL9 (See Table S7). Visual inspection in Bandage showed highly fragmented assemblies for these two isolates.

Both of these isolates were fluconazole-resistant without another established mutational mechanism. Mitochondrial deletions leading to petite phenotype are known to cause fluconazole resistance in *N. glabratus*, via *PDR1*-dependent mechanisms (14). Both cases were preceded by triazole exposure (20 days of fluconazole treatment in an intraabdominal infection, and 15 days of posaconazole prophylaxis in neutropenia, respectively). Both demonstrated a petite phenotype (see Figure S2). Isolate AUH_NAGL1 occurred concomitantly with a grande (large) colony type which was fluconazole-susceptible. The respiration-deficient phenotype in these petite variants was confirmed by failure to grow on a non-fermentable carbohydrate medium (glycerol agar) (Figure S3).

### Pair analysis

Genes with mutations restricted to the triazole-resistant partners of three or more phylogenetic pairs included *PDR1* and seven genes with predicted functions including GPI-anchored cell wall proteins and adenylate cyclase. Further details are included in Table S9.

## DISCUSSION

In our seven-year study of *N. glabratus* isolates at a quarternary hospital, we observed a largely polyclonal *N. glabratus* population structure. Antifungal resistance was predominantly polyphyletic, suggesting independent acquisitions, but a small cluster with the same resistance mechanism was observed. In the ST301 cluster, the common factor of ICU admission and a universal *PDR1* mutation suggests clonal fAMR transmission across five years. The pairwise SNP distances defining this cluster (256-417 SNPs) are compatible with transmission, noting previous reports of distances of 2-936 SNPs within *N. glabratus* clusters (34), and distances of 194-451 SNPs (4), and approximately 1000 SNPs (5) within *N. glabratus* representing probable transmission.

Novel findings included putative loss-of-function mutations in *PDR1* in a hypersusceptible isolate (fluconazole MIC 1 μg/ml), and in *FCY2* in an isolate with a very high flucytosine MIC (32 μg/ml). *PDR1* disruption is known to increase fluconazole susceptibility (11). *FCY2* deletions have been associated with *in vitro* flucytosine nonsusceptibility in *Saccharomyces cerevisiae* (35) and *Clavispora lusitaniae* (36). The role of *FCY21* in flucytosine nonsusceptibility is uncertain, but the finding of stop and missense mutations in three Non-susceptible isolates warrants future confirmation. *In vitro* disruption of *FCY21* in *S. cerevisiae* is associated with reduced flucytosine susceptibility (37).

Missense mutations in efflux pump-encoding genes were observed in several triazole-non-susceptible isolates; their significance remains to be determined. Efflux pump overexpression, for example via *PDR1* gain of function, is more established as a resistance mechanism than coding sequence mutations (23). The *ERG3* mutation H218R, occurring only in the amphotericin B-Non-susceptible isolate, might theoretically contribute to polyene resistance through reduced ergosterol production, but functional confirmation is required.

Structural variation was noted in three non-susceptible isolates. In one isolate, high-level triazole resistance might be explained by the combination of *ERG11* CNV, an *ERG11* mutation, and *PDR1* gain-of-function. Although comparison with an international dataset suggests Chromosome E duplication might not be sufficient in isolation, it has previously been described in treatment-emergent resistance (12). In two isolates, apparent deletions within the mitochondrial genome represent a likely mechanism of resistance. In both cases, prior triazole exposure might have selected for mitochondrial genome loss leading to a respiratory-deficient phenotype capable of surviving under triazole stress. Inclusion of methods for detecting mitochondrial genome deletions should be considered in future bioinformatic pipeline development.

Fungal antimicrobial resistance is complex and significant gaps in our understanding remain (38). The lack of identified resistance mechanisms in 10 isolates might reflect (i) transient mechanisms, such as altered efflux regulation or unstable aneuploidies; (ii) colony selection in a heteroresistant infection; or (iii) a mutational mechanism outside the genes assessed. In the isolate showing low-level anidulafungin resistance, the mutations in *FKS1, FKS2* and *FKS3* are of uncertain significance: they were present in Susceptible isolates; they are outside resistance hot spots; and previously reports have not consistently associated these with echinocandin resistance (39, 40). A stress-induced compensatory increase in chitin production is a possible alternative mechanism of resistance (41).

Our findings should be interpreted with caution, with selection bias inherent in use of stored isolates, a small sample size and the single-centre retrospective design. Since our definition of Non-susceptibility prioritised specificity in the case of triazoles other than fluconazole, we may have excluded some mutational mechanisms contributing to low-level triazole non-susceptibility.

Our study adds to a growing body of literature on hospital-acquired *N. glabratus* infections by considering structural variation in addition to mutational analysis. Our inclusion of hospital epidemiological data contributes to hypotheses regarding fAMR transmission dynamics. Sporadic nosocomial transmission has been reported, but evidence of true transmitted resistance has been limited. Our reporting of a shared *PDR1* SNP in a monophyletic cluster supports nosocomial transmission of fAMR. A low representation of echinocandin resistance is a limitation reflective of our local context (5, 42).

Future studies systematically and longitudinally capturing commensal and invasive isolates would aid our understanding of nosocomial transmission of fAMR. Assessing within-host and within-cluster diversity would enhance the accuracy of phylogenetic interpretation. Future study should also confirm the suspected *ERG11* CNV and mitochondrial deletions. Other research priorities include the contribution of epigenetics and post-translational modifications, to define further the resistance mechanisms relevant to clinical isolates and patient management for this important and emerging drug-resistant pathogen.

## ACKNOWLEDGEMENTS

The authors received no specific funding for this study. BPH, NLS and AT are supported by Investigator Grants from the National Health and Medical Research Council Australia (GNT1196103 to BPH, GNT2033803 to NLS and 2033452 to AT). GR is supported by a NMHRC PhD Scholarship (#2013970). The authors declare no conflict of interest.

The authors would like to acknowledge with gratitude the Austin Pathology Department of Microbiology as the source and owner of the clinical isolates. The authors would like to acknowledge the contributions of many people to this study including the Austin Pathology Department of Microbiology, Melbourne, particularly Elizabeth Grabsch, Kit Liu, Tirzah Korrapadu, Jenny Wang and Tram Nguyen; Taylor Harshegyi (Centre for Pathogen Genomics Innovation Hub, Melbourne); Catherine Glover and Sandra Johnston (Epidemiology Section, Microbiological Diagnostic Unit Public Health Laboratory, Melbourne); Himal Shrestha (Bioinformatics Section, Microbiological Diagnostic Unit Public Health Laboratory, Melbourne); Celeste Donato (Australian Pathogen Genomics Program, Melbourne); Ryan Wick (Department of Microbiology and Immunology, University of Melbourne); Ammar Aziz (Victorian Infectious Diseases Reference Laboratory, Melbourne); and Cameron Patrick and Jeremy Metha (University of Melbourne).

## AUTHOR CONTRIBUTION STATEMENT

AG-W, JK, NS, BH and TS conceptualised and designed the study. JK, NS, BH and TS supervised the project. ML provided the isolates and contributed to the study design. GR assisted with the study design and ethics application. AT provided technical advice and feedback on the manuscript. AG-W performed the isolate retrieval, susceptibility testing, medical record review, statistical analysis, figure generation and draft manuscript writing. LJ performed the sequencing with AG-W and contributed to the study design. AG-W, TS, KH and JL performed the bioinformatic analysis. DD and MDA assisted with the wet laboratory work. SV provided expertise for the statistical analysis. RG and SG provided input into the bioinformatic analysis and manuscript. All authors reviewed the manuscript.

## DATA AVAILABILITY

Sequence reads are deposited in the National Center for Biotechnology Information Sequence Read Archive (Bioproject accession: PRJNA856410. Bioinformatic analysis scripts are available from the project github repository (github.com/andrewgw1989/Nakaseomyces_glabratus_genomics). Sample metadata are provided in the Supplemental Material.

## ETHICS APPROVAL

Approval was received from the Austin Health Human Research Ethics Committee (reference: HREC/92327/Austin-2023).

